# Novel Spike-stabilized trimers with improved production protect K18-hACE2 mice and golden Syrian hamsters from the highly pathogenic SARS-CoV-2 Beta variant

**DOI:** 10.1101/2023.07.07.548077

**Authors:** Carlos Ávila-Nieto, Júlia Vergara-Alert, Pep Amengual-Rigo, Erola Ainsua-Enrich, Marco Brustolin, María Luisa Rodríguez de la Concepción, Núria Pedreño-Lopez, Jordi Rodon, Victor Urrea, Edwards Pradenas, Silvia Marfil, Ester Ballana, Eva Riveira-Muñoz, Mònica Pérez, Núria Roca, Ferran Tarrés-Freixas, Julieta Carabelli, Guillermo Cantero, Anna Pons-Grífols, Carla Rovirosa, Carmen Aguilar-Gurrieri, Raquel Ortiz, Ana Barajas, Benjamin Trinité, Rosalba Lepore, Jordana Muñoz-Basagoiti, Daniel Perez-Zsolt, Nuria Izquierdo-Useros, Alfonso Valencia, Julià Blanco, Bonaventura Clotet, Victor Guallar, Joaquim Segalés, Jorge Carrillo

## Abstract

Most COVID-19 vaccines are based on the SARS-CoV-2 Spike glycoprotein (S) or their subunits. However, the S shows some structural instability that limits its immunogenicity and production, hampering the development of recombinant S-based vaccines. The introduction of the K986P and V987P (S-2P) mutations increases the production of the recombinant S trimer and, more importantly, its immunogenicity, suggesting that these two parameters are related. However, S-2P still shows some molecular instability and it is produced with low yield. Thus, S-2P production can be further optimized. Here we described a novel set of mutations identified by molecular modelling and located in the S2 region of the Spike that increase S-2P production up to five-fold. Besides their immunogenicity, the efficacy of two representative S-2P-based mutants, S-29 and S-21, protecting from a heterologous SARS-CoV-2 Beta variant challenge was assayed in K18-hACE2 mice (an animal model of severe SARS-CoV-2 disease) and golden Syrian hamsters (GSH) (a moderate disease model). S-21 induced higher level of WH1 and Delta variants neutralizing antibodies than S-2P in K18-hACE2 mice three days after challenge. Viral load in nasal turbinate and oropharyngeal samples were reduced in S-21 and S-29 vaccinated mice. Despite that, only the S-29 protein protected 100% of K18-hACE2 mice from severe disease. When GSH were analyzed, all immunized animals were protected from disease development irrespectively of the immunogen they received. Therefore, the higher yield of S-29, as well as its improved immunogenicity and efficacy protecting from the highly pathogenic SARS-CoV-2 Beta variant, pinpoint the S-29 spike mutant as an alternative to the S-2P protein for future SARS-CoV-2 vaccine development.

**Authors summary:** The rapid development of SARS-CoV-2 vaccines have been pivotal in the control of the COVID-19 pandemic worldwide. Most of these vaccines include the S glycoprotein as the main immunogen since this protein, and particularly its receptor binding domain (RBD), is the major target of neutralizing antibodies. SARS-CoV-2 have been evolving from the beginning of the pandemic and several variants with increased transmissibility, pathogenicity or resistance to infection– or vaccine-induced immunity have emerged. Different strategies have been adopted to improve vaccine protection including additional booster doses or the adaptation of the S immunogens to the novel SARS-CoV-2 variants. As a complementary strategy we have identified a combination of non-proline mutations that increase S production by 5-fold (S-29 protein). Despite the sequence of this novel S-29 immunogen is based on the ancestral SARS-CoV-2 WH1 variant, it effectively protects animal model from the highly pathogenic and neutralization resistant SARS-CoV-2 Beta variant. Thus, we describe a novel set of mutations that can increase the production and efficacy of S-based COVID-19 vaccines.

## Introduction

Vaccines have been extensively used to control infectious diseases. While smallpox is the only pathogen that has been eradicated in human, mass immunization programs have reduced the spread of other infectious diseases, including tetanus, polio, and measles[1,2]. In January 2020, the severe acute respiratory syndrome coronavirus 2 (SARS-CoV-2) was identified as the causal agent of the coronavirus disease 2019 (COVID-19), and many laboratories rapidly started programs to develop COVID-19 vaccines[3,4]. To date, several COVID-19 vaccines are available and have contributed to the reduction of COVID-19 impact on public health[5,6]. However, COVID-19 is still present in the world and new SARS-CoV-2 variants continue emerging with high transmissibility and/or resistance to the immune responses elicited after infection and/or vaccination[7].

Within the SARS-CoV-2 proteome, the Spike (S) mediates virus attachment by binding to the angiotensin-converting enzyme 2 receptor (ACE2) on the surface of target cells. After being primed by host proteases, S promotes viral entry and cell infection[8,9]. Therefore, most SARS-CoV-2 vaccines are based on this protein since it is the main target of neutralizing antibodies (NAbs) [particularly, the receptor binding domain (RBD)][3,4,10]. The S is a trimer, and each monomer has two subunits: the S1 extracellular, and the S2 membrane anchor subunits. While the S1 binds to ACE2 via the RBDs, the S2 domain participates in the membrane fusion process, which involves drastic conformational changes[8,11,12]. Thus, the S glycoprotein shows a certain degree of structural instability that might hamper its production as recombinant protein and modulate its immunogenicity. This feature is shared with functional homologous surface proteins from other viruses, including the S of Middle East respiratory syndrome coronavirus (MERS-CoV), the Fusion protein of the respiratory syncytial virus (RSV), or the Envelope glycoprotein of the human immunodeficiency virus (HIV)[13]. Several studies have shown that it is possible to stabilize these proteins in their prefusion state and improve their production and immunogenicity[13]. In this sense, the introduction of two proline mutations in the S2 (S-2P) has been proposed as a common strategy for the stabilization of this glycoprotein from several coronaviruses, including the SARS-CoV-2 [11,14]. In fact, some of the most used SARS-CoV-2 vaccines, such as BNT162b2, mRNA-1273 or Ad26.COV2.S are based on the S-2P strategy [15–17].

Importantly, the S-2P protein still retains some structural instability and generates a poor yield when the protein is produced as recombinant protein (around 0.5 mg/L)[11,18]. Several studies have addressed these limitations by introducing additional stabilizing mutations. In this sense, the incorporation of four additional prolines (S-6P) improved the stability of the S trimer and increased its yield by ten-fold[18]. In another approach, the incorporation of the mutations D614N, A892P, A942P, and V987P stabilized the S protein in a close-prefusion state and increased its yield by 6-fold[19]. However, whereas the addition of disulfide bridges between different domains of the S glycoprotein reduced the motility of the RBD, it failed improving recombinant trimer production[20,21]. Alternatively, pre-fusion stabilizing mutations have also been identified by high-throughput methods. Thus, the addition of D994Q mutation to the S-2P backbone increased its production as soluble recombinant protein by three-fold[22]. However, it remains poorly understood how these mutations modify the S immunogenicity compared to S-2P.

Here, we describe a set of novel mutations that increase SARS-CoV-2 S yields by five-fold, while maintaining the immunogenicity and protection efficacy against the development of SARS-CoV-2-induced disease in K18-hACE2 transgenic mice and golden Syrian hamsters (GSH) previously observed with the S-2P prototype.

## Results

### Strategy for S glycoprotein stabilization

To increase the S glycoprotein stability and immunogenicity, we followed two different approaches: 1) introduction of point mutations into the S sequence to increase its stabilization (using the open state as a reference structure), and 2) increase of RBD exposure by forcing an open conformation. In this regard, we built a computational pipeline involving the three-dimensional modeling of all possible single mutations in both scenarios (see the Methods section for more details). Moreover, all single mutations that showed a preference for any of these two conditions were visually inspected. Mutations that clearly generated well-defined interactions (e.g. hydrogen bonds, ionic interactions for filling hydrophobic pockets) between different chains of the S trimer were prioritized (Fig. 1A).

**Fig. 1.**
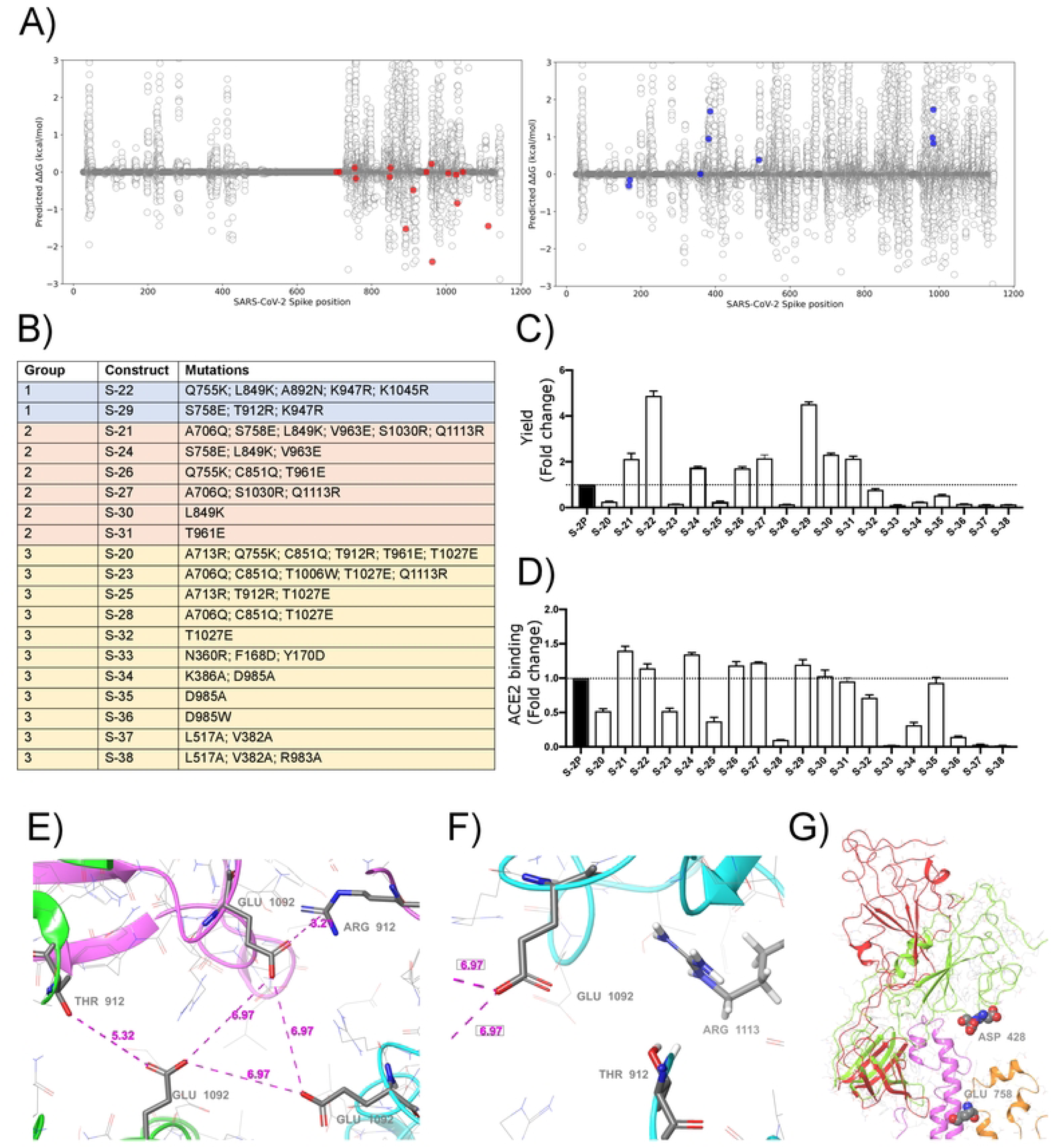
Selection of mutations that stabilized S glycoproteins. **A.** S stabilizing mutations (red variants, left plot), or amino acids changes that increased RBD exposition (blue variants, right plot) were selected based on energetic filters and visual inspection. Positive energy values indicate stabilization of the open structure versus the closed one. Mutations with neutral (or slightly opposite) energetic trend were included. **B** List of S constructs that incorporate the selected mutations identified in a. **C** Yields of recombinant S mutants in a five-day cell culture supernatant. Mean plus standard deviation of three experiments are shown. **D** RBD exposure index in selected recombinant proteins. Data are shown as ratio between RBD binding and total protein. Mean plus standard deviation of three experiments are shown. **E** Presence of a cluster of three Glu residues (one Glu1092 from each chain) that are facing each other in close proximity, with no positive residues nearby. Location of Thr912 is underlined as well. Also notice that one of the three Thr has been mutated to Arg clearly showing a salt bridge interaction with the glutamic acid. **F** A detail of the proximity of the 1113 residue, already mutated to Arg, to the Glu1092 cluster. Thr912 is also showed. Structure models were based on 6VXX PDB structure. **G** Detail of the RBD opening process and location of some key residues. The red and green ribbons indicate the difference between the open and closed states, underlying the position of the two consecutive aspartic acid residues, Asp428 and Asp427, at the tip of the RBD domain. In orange ribbons the location of the S758E mutation is shown. Notice that the inserted glutamic residue collides with the neighbor helix (pink ribbons). Models generated from the 6VXX (closed) and 6VYB (open) PDB structures.

S mutants (Fig. 1B) were then produced, and yields evaluated (Fig. 1C). Based on their production levels, these glycoproteins were classified into three different groups (Fig. 1B and C). Group 1 included those constructs (i.e., S-29 and S-22) that were produced at the highest levels (five-fold compared to the S-2P protein). Group 2 contained S-21, S-24, S-26, S-27, S-30, and S-31, whose production was intermediate (two-fold higher than S-2P). Last, Group 3 included those S mutants with a protein yield lower than S-2P (S-20, S-23, S-25, S-28, S-32, S-33, S-34, S-35, S-36, S-37, and S-38). Remarkably, all constructs designed to increase RBD exposure were in Group 3, suggesting that those mutations drastically impacted the S stability and/or its production. However, most constructs with an improved production, also showed a better RBD exposure (Fig. 1C and D).

Variants S-22 and S-29, with higher expression yield, introduced a positive charge per chain in a local area where the Glu1092 of each chain might cause destabilization. Figure 1E shows the presence of this cluster of Glutamic acid residues facing each other and how the T912R mutation in S-29 might introduce significant stabilization. Similarly, the Q1113R mutant in S-22 placed an arginine next to Glu1092 (Fig. 1F). Analogous observations can be extracted of most mutants introducing a net charge. We also observed that most mutants increasing RBD exposure, such as S-21, S-24 and S-29, incorporated the S758E mutation. This mutation is in the vicinity of the tip of the closed RBD domain, where two consecutive Aspartic acid residues, Asp427 and Asp428 are located (Fig. 1G). After modeling the possible positioning of Glu758 (with an initial significant clash with a helix backbone), we speculate that it would be displaced towards the tip of the RBD domain and destabilize the closed conformation by electrostatic repulsion.

Interestingly, S-22 and S-29 constructs share the conservative mutation K947R located in the middle of the heptad repeat 1 (HR1) helix, which could enhance the thermal stability of the protein[23].

### S-21 and S-29 vaccination protects K18-hACE2 mice from SARS-CoV-2-induced disease

To investigate the impact of S mutations on its immunogenicity and capability to protect from SARS-CoV-2-induced disease, we selected two representative S mutants from group 1 (S-29) and 2 (S-21). Then, we performed an immunization study using K18-hACE2 transgenic mice that were subsequently challenged with SARS-CoV-2 B.1.351 (Beta) variant (Fig 2A). We used this experimental design for the following reasons: 1) K18-hACE2 transgenic mice develop a severe form of the disease that leads to death[24] unless animals are vaccine-protected; 2) the SARS-CoV-2 Beta variant is partially resistant to antibodies elicited by natural infection or vaccination with immunogens based on the original strain (Wuhan, WH1)[25]; and 3) the SARS-CoV-2 Beta variant is one of the most pathogenic SARS-CoV-2 variants tested in K18-hACE2 transgenic mice[24]. Thus, we established five experimental groups: S-2P (n=21), S-21 (n=22), S-29 (n=22), infected positive controls (n=16), and uninfected negative controls (n=10). Mice from S-2P, S-21, and S-29 groups were immunized twice, three weeks apart. Animals from both control groups received antigen-free doses. Two weeks after the boost, all animals (except the negative controls) were intranasally challenged with the SARS-CoV-2 Beta variant. Blood and tissue samples were collected after viral challenge on days 3 (n=6), 6 (n=6) and 14 (n=10 for S-21, S-29 and uninfected controls, and n=8 for S-2P) to analyze tissue damage and viral replication (Fig. 2A). All mice that developed severe disease (one mouse in the S-2P and S-21 groups, and all mice from the positive control group) were euthanized before day 14 following humane endpoints and were analyzed separately.

**Fig. 2.**
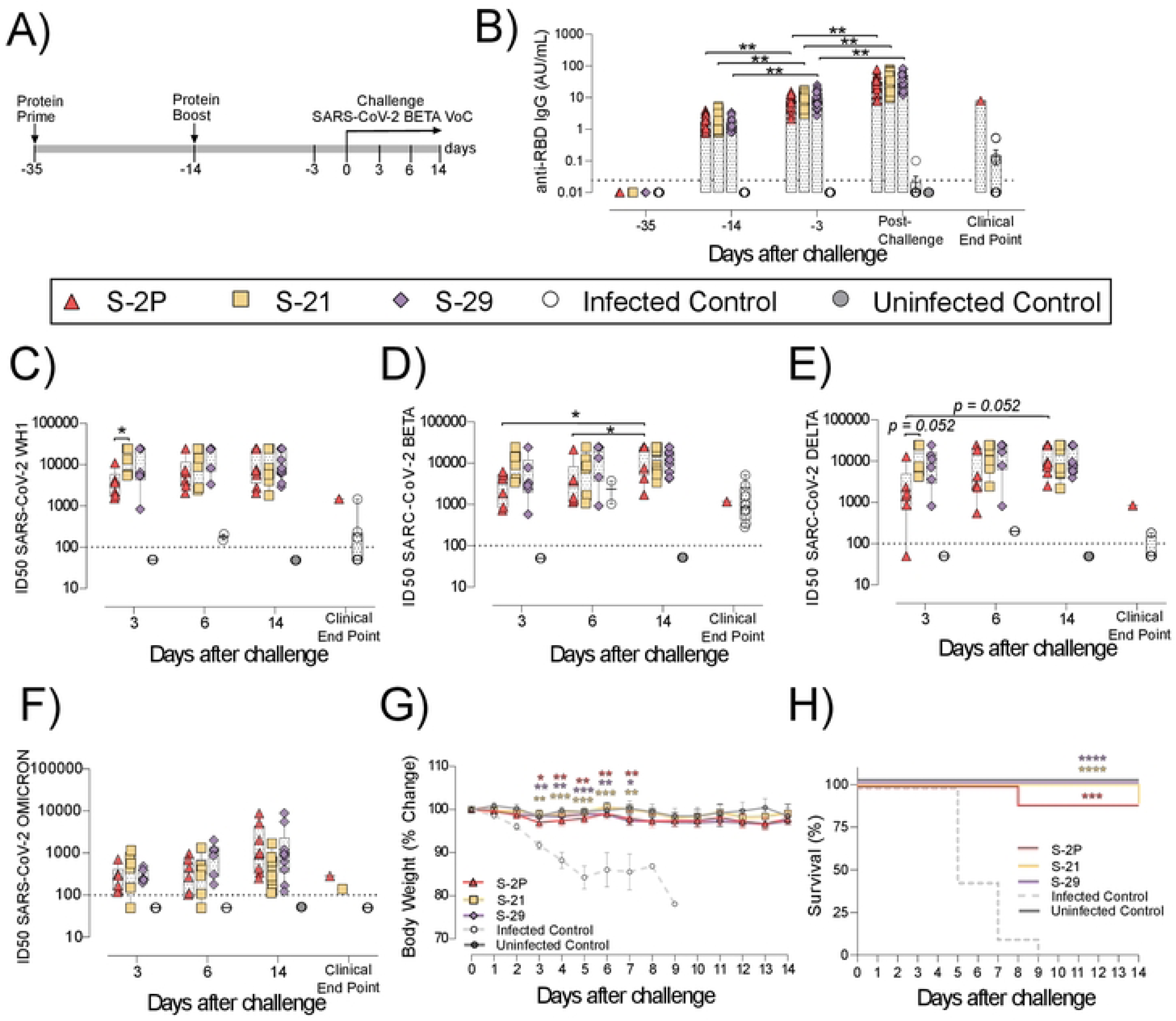
Prophylactic activity of S-21 and S-29 immunization and vaccine-induced humoral response elicited in K18-hACE2 transgenic mice challenged with SARS-CoV-2 B.1.351 (Beta) variant. K18-hACE2 transgenic mice were immunized following a prime/boost strategy with S-21, S-29, or S-2P, and challenged with SARS-CoV-2 Beta. The vaccine-induced humoral responses, weight changes, and survival of mice were evaluated after immunization and/or viral challenge. **A** Overview of immunization strategy and infection timeline. Biological samples were collected at indicated time points. **B** Kinetics of anti-RBD antibodies in serum samples. Red triangles: S-2P group (n=21). Yellow squares: S-21 (n=22). Purple diamond: S-29 (n=22). White circles: unvaccinated-challenge mice (n=16). Grey circles: unvaccinated-uninfected mice (n=10). Groups in each time point were analyzed using Kruskal-Wallis and Conover’s post-hoc tests with multiple comparison correction by FDR. Differences among animals within a particular group along time were analyzed using Friedman and Conover’s post-hoc test for paired data with FDR correction. Sera neutralizing activity against: **c** SARS-CoV-2 WH-1 variant, **D** B.1.351 (Beta) variant, **e** B.1.617.2 (Delta), and **f** B.1.1.529 (Omicron) variants after viral challenge. Neutralization data were analyzed as indicated in “b”. **G** Percentage of weight variation in SARS-CoV-2 B.1.351 infected K18-ACE2 mice over time. Statistical analysis was performed against the unvaccinated and challenged group using Kruskal Wallis with Dunńs post-hoc test. **H** Kaplan-Meier plot showing the percentage of SARS-CoV-2-infected animals that were disease-free at the end of the experiment. Statistical analysis was performed against unvaccinated group using Mantel-Cox test. * *p*<0.05, ** *p*<0.01, *** *p*<0.001, **** *p*<0.0001. Mean plus standard errors of the means (SEM) are shown.

Anti-RBD (Fig. 2B) and anti-S (S1A Fig) IgG humoral responses were evaluated prior to each immunization and viral challenge, and in euthanized animals after infection on days 3, 6, and 14, or due to humane endpoints. Regardless of the immunogen used, all vaccinated animals developed similar anti-RBD (Fig. 2B) and anti-S IgG levels (S1A Fig), which increased after each immunization and after viral challenge (p<0.01, Conoveŕs post-hoc test). Since we did not identify significant differences in the humoral responses among vaccinated groups after challenge (S1B and C Fig), we pooled these mice in a single “post-challenge” group to simplify the analysis. Of note, unvaccinated but challenged positive controls elicited low levels of anti-S and anti-RBD IgG antibodies (Fig. 2B and S1A Fig) that were detected from day 6 after viral challenge (S1B and 1C Fig).

Sera neutralizing activity against SARS-CoV-2 WH1, Beta, Delta, and Omicron variants was detected in all three vaccinated groups (Fig. 2C-F). Interestingly, despite having similar levels of anti-RBD IgGs (Fig. 2B), and a slightly higher levels of anti-S IgG antibodies (S1A Fig), S-2P vaccinated mice showed lower sera neutralizing activity against WH1 and Delta variants on day 3 than those immunized with S-21 (Fig 2C and E) (WH1 p<0.05; Delta p=0.052, Conoveŕs post-hoc test). Sera neutralizing activity against Delta and Beta variants increased over time in the S-2P vaccinated group after viral challenge (Beta p<0.05; Delta p=0.052, Conoveŕs post-hoc test), suggesting that infection boosted the humoral response in these animals. In line, unvaccinated mice developed low sera neutralizing activity against the SARS-CoV-2 Beta variant (Fig 2D) with some cross-reactivity with WH1 but limited cross-neutralizing activity against other SARS-CoV-2 variants (Fig. 2C, 2E, and 2F) after viral challenge. No boost effect on sera neutralizing activity was detected in S-21– and S-29-immunized mice after challenge, suggesting that the humoral response reached a plateau in these groups.

To determine whether S-2P-, S-21-, and S-29-vaccinated mice were protected against SARS-CoV-2-induced severe disease, we measured body weight evolution (Fig. 2G), clinical sings, and survival rate after viral challenge (Fig. 2H). A progressive weight loss was observed in all unvaccinated but challenged mice starting on day 2 post-challenge. These mice developed a severe disease on days 5-9 post-infection and were euthanized following humane endpoints. Conversely, all vaccinated mice (except one S-2P– and one S-21-immunized mice), were disease-free (Fig. 2H) and did not experience weight loss. All mice belonging to S-29 group were protected from severe disease development (Fig. 2G and 2H).

The presence of SARS-CoV-2 in oropharyngeal swabs and tissue samples from nasal turbinate, lung, and brain was analyzed by RT-qPCR. All vaccinated mice had significantly lower levels of genomic viral RNA (gRNA) in lung on day 3 post-inoculation compared to the positive control group (p<0.05, Peto & Peto Left-censored k-sample test) (Fig. 3A). Most notably gRNA was scarcely detected in brain of vaccinated mice compared to unvaccinated animals (Fig. 3A). Interestingly, S-21– and S-29-vaccinated mice showed lower viral load in nasal turbinate than S-2P and control groups on day 3, and S-2P vaccinated mice on day 6 (p<0.05, Peto & Peto Left-censored k-sample test) (Fig. 3A). The lack of differences with the control group on day 6 could be explained due to the small number of unvaccinated mice that reached this timepoint, since the majority had been euthanized on day 5 post-infection (Fig 2H). Similarly, S-21 and S-29 groups exhibited lower viral loads in oropharyngeal swabs than unvaccinated mice (S-21 p<0.05; S-29 p=0.066; Peto & Peto Left-censored k-sample test) (Fig. 3A). Generally, gRNA decreased over time in all immunized mice regardless of the analyzed sample, whereas the opposite outcome was observed in mice belonging to the challenged control group, and in those vaccinated mice that developed severe disease (Fig. 3A). Similar results were observed when subgenomic viral RNA (sgRNA) was analyzed using the same biological samples (S1D Fig).

**Fig. 3.**
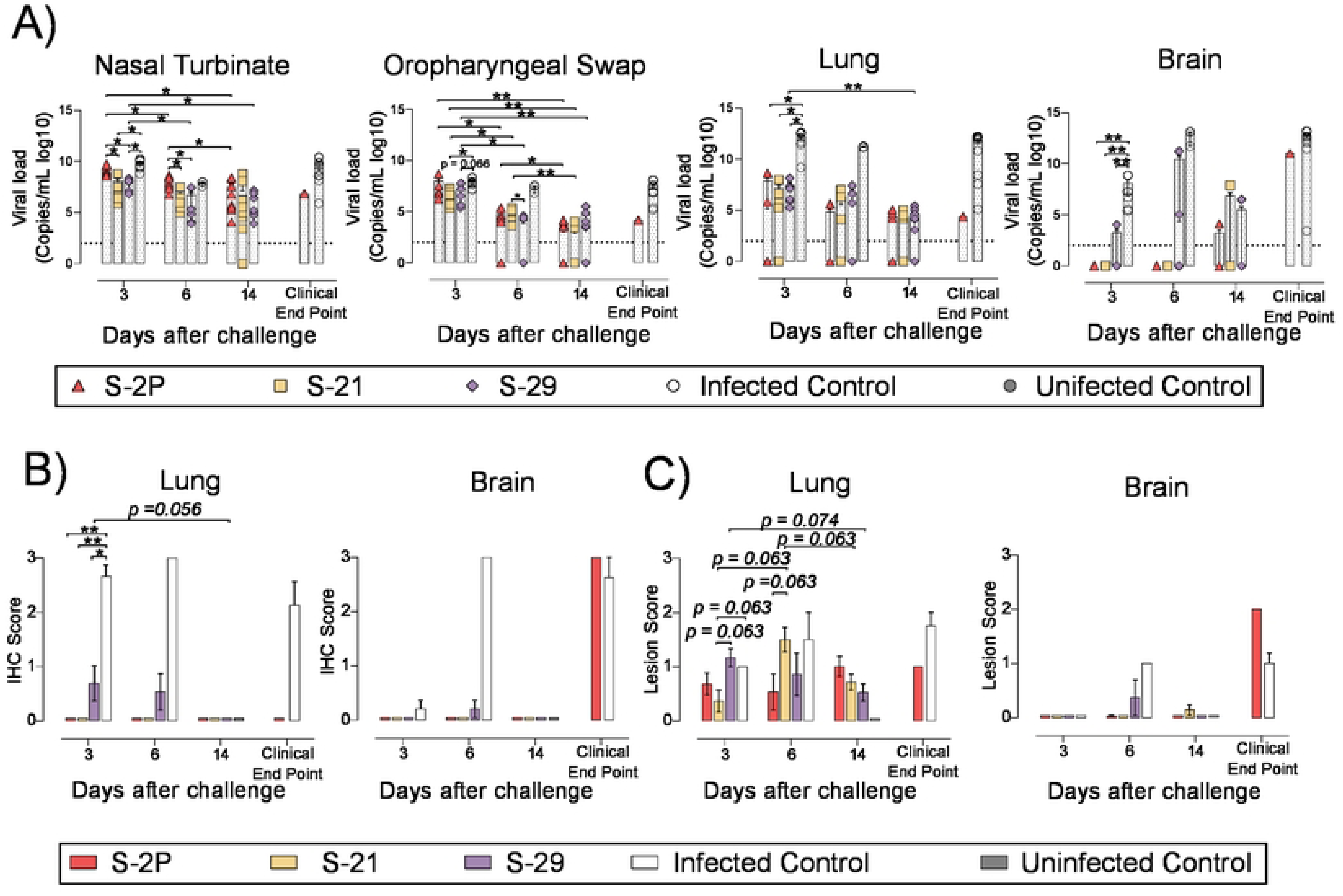
Viral load and histopathology analysis of biological samples from SARS-CoV-2 B.1.351 infected K18-hACE2 transgenic mice after vaccination. SARS-CoV-2 viral loads were analyzed in oropharyngeal swabs, and samples from nasal turbinate, lung, and brain of infected K18-hACE2 mice. Virus distribution and tissue damage were analyzed by immunohistochemistry and histopathology, respectively. **A** Levels of SARS-CoV-2 gRNA (expressed as logarithmic of copies/mL) in oropharyngeal swabs, nasal turbinate, lung, and brain during infection. Dotted line indicates limit of detection (100 copies/mL). Differences among groups were analyzed using Peto & Peto left-censored k-sample test with FDR correction. **B** Histopathological analysis of brain, lung, and nasal turbinate by hematoxylin and eosin staining. Lesion score: (0) no lesion, (1) mild lesion, (2) moderate lesion, and (3) severe lesion. **C** Detection of SARS-CoV-2 nucleocapsid protein in brain, lung, and nasal turbinate by immunohistochemistry. Staining score: (0) no antigen, (1) low, (2) moderate, and (3) high viral antigen. Differences among groups were analyzed using Asymptotic Generalized Pearson Chi-Squared test with FDR correction. * *p*<0.05, ** *p*<0.01.

To confirm active viral replication, nucleoprotein (NP) levels were analyzed by immunohistochemistry (IHC). NP was hardly detected in lung and brain samples from S-2P, S-21 and S-29 groups (Fig. 3B). These data are in accordance with the viral loads detected in these samples. Despite that, some tissue damage was still detected in the lungs of all vaccinated groups. Remarkably, limited tissue damage was found in the brain of vaccinated mice, except in those animals euthanized due to humane endpoints (Fig. 3C).

Overall, S-2P, S-21, and S-29 trimers displayed an equivalent immunogenicity in K18-hACE2 transgenic mice and protected these animals from developing severe disease after SARS-CoV-2 Beta variant challenge. Interestingly, S-21– and S-29-immunized animals had lower viral loads in nasal turbinate than S-2P and infected controls on days 3 and 6 after challenge. Viral loads were also reduced in oropharyngeal swabs of these mice on day 3 compared to infected control groups.

### S-21 and S-29 trimer vaccination protects golden Syrian hamsters from COVID-19 development

To confirm the results obtained in K18-hACE2 mice, a second immunization and challenge experiment was performed using GSH with the same immunogens. Unlike K18-hACE2 mice, GSH develop a moderate form of SARS-CoV-2-induced disease, from which they spontaneously recover by day 14 after challenge[26,27]. GSH were immunized following a similar prime/boost strategy to the previously used for K18-hACE2 transgenic mice. Animals were intranasally challenged with the SARS-CoV-2 Beta variant and followed up until day 7 post-challenge (Fig. 4A), since it has been described that GSH start recovering weight from this day[26,27].

**Fig. 4.**
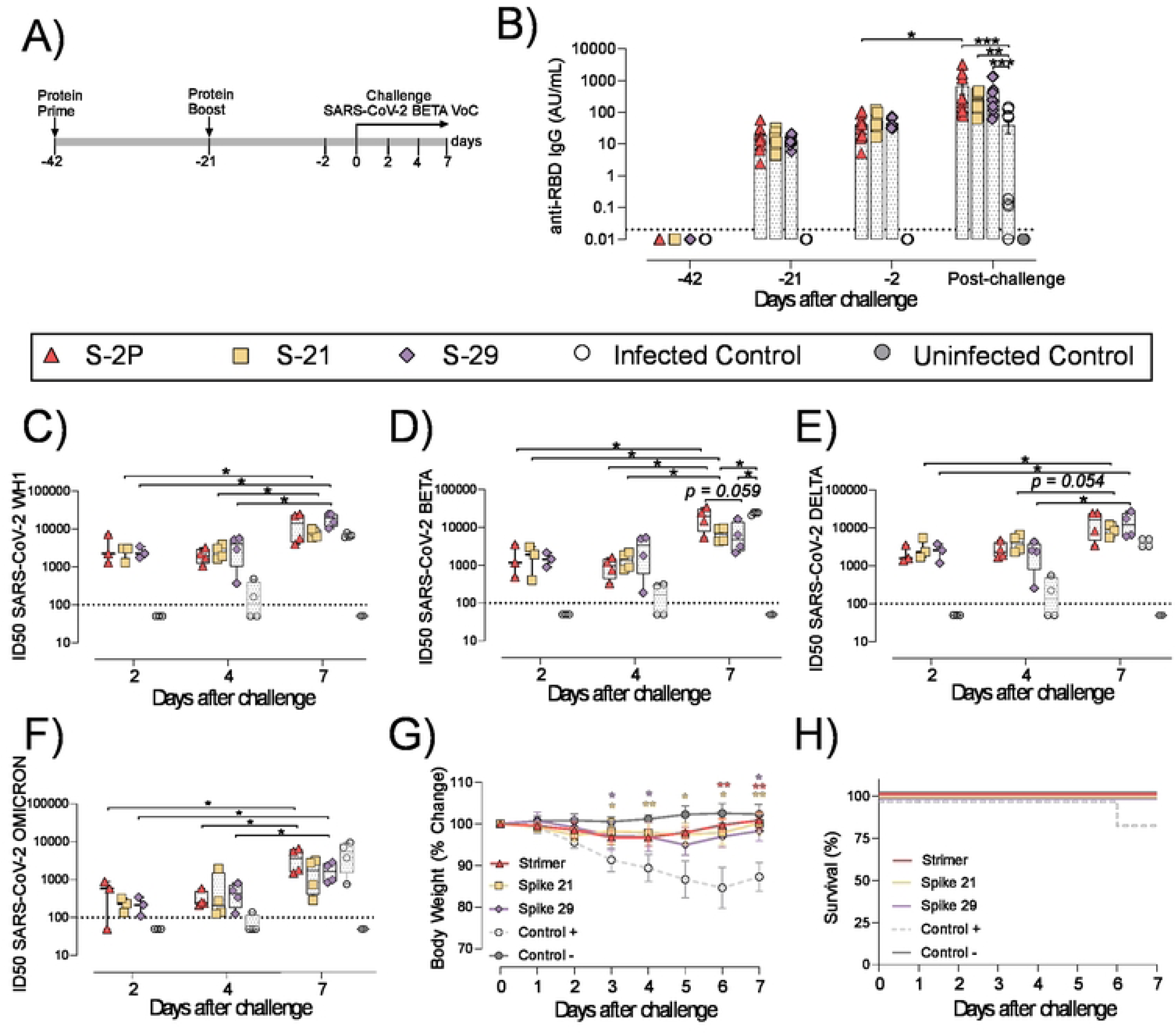
Vaccine-induced humoral responses and prophylactic activity of S-21 and S-29 in immunized GSH after challenge with the SARS-CoV-2 B.1.351 (Beta) variant. GSH were immunized twice with S-21, S-29 or S-2P, and subsequently challenged with SARS-CoV-2 B.1.351 variants. The humoral response, weight changes, and survival of mice were evaluated after immunization and/or viral challenge. **A** Outline of immunization schedule and infection timeline. Biological samples were collected at the indicated time points. **B** Kinetics of anti-RBD antibodies in serum samples. Red triangles: S-2P group (n= 11). Yellow squares: S-21 (n=11). Purple diamond: S-29 (n=11). White circles: unvaccinated-challenged mice (n=11). Grey circles: unvaccinated-uninfected mice (n=5). Groups in each time point were analyzed using Kruskal-Wallis and Conover’s post-hoc tests with multiple comparison correction by FDR. Differences among animals within a particular group along time were analyzed using the Friedman and Conover’s post-hoc tests for paired data with FDR correction. Sera neutralizing activity after viral challenge against: **C** SARS-CoV-2 WH-1, **D** B.1.351 (Beta), **E** B.1.617.2 (Delta), and **F** B.1.1.529 (Omicron) variants. Neutralization data were analyzed as indicated in **B**. **G** Percentage of weight variation in SARS-CoV-2 B.1.351-infected GSH over time. **H** Kaplan-Meier plot showing the frequency of disease-free SARS-CoV-2-infected animals at the end of the experiment. Statistical analysis was performed against the unvaccinated group using Kruskal Wallis and Dunńs post-hoc tests. * p<0.05, ** p<0.01, *** p<0.001.

In accordance with K18-hACE2 transgenic mice data, the three vaccinated groups (S-2P, S-21 and S-29) developed similar levels of anti-RBD and anti-S binding IgG (Fig. 4B, S2A and S2B Fig). Interestingly, the second immunization did not boost vaccine-induced anti-S or anti-RBD IgG antibodies, indicating that a second vaccine dose might not be needed in this animal model. Interestingly, an increase in anti-S IgG levels was observed after viral challenge in both vaccinated and unvaccinated but challenged mice (S2A Fig). However, when anti-RBD IgG responses were analyzed, that boosting effect was less evident and only detected in the S-2P and in unvaccinated and challenged groups (Fig. 2B and S2B Fig). These results suggest that viral challenge elicited a rapid humoral response in naïve animals, boosting anti-S IgG responses, but had little effect in vaccinated GSH. Despite that, immunized GSH showed higher levels of anti-S and anti-RBD antibodies than challenged controls (p<0.001 for S-2P and S-29 group, and p<0.01 for S-21 group; Friedman test) (Fig. 4B and S2A Fig).

Sera neutralizing activity against SARS-CoV-2 WH1, Beta and Delta, and to a lesser extent, Omicron variants was detected in vaccinated animals at all post-challenge timepoints (Fig. 4C-F). No differences were identified among immunized groups. Remarkably, and contrarily to K18-hACE2 transgenic mice vaccine study data, sera neutralizing activity against all four SARS-CoV-2 variants were observed in some challenged positive control animals by day 4 after challenge (Fig. 4C-F). These results indicate that cross-reactive neutralizing antibodies were generated in those individuals. Unexpectedly, the neutralizing activity against the SARS-CoV-2 Beta variant was higher in challenged control animals than in S-21– and S-29-vaccinated GSH by day 7 (Fig. 4D). According to the binding ELISA data, neutralization titers also increased in immunized GSH by day 7 after viral challenge (p<0.05; Conoveŕs post-hoc test), indicating that infection boosts vaccine-induced humoral neutralizing responses (Fig. 4C-F).

To determine whether vaccination protected GSHs from SARS-CoV-2-induced disease, we monitored animal weight over time after viral challenge (Fig. 4G). Challenged control GSHs showed progressive weight reduction until day 6, which was indicative of disease progression. One animal from this group suffered a weight reduction greater than 20% by day 6 post-inoculation and was euthanized according to humane endpoints (Fig. 4H). No significant weight loss was observed in vaccinated GSHs, indicating that these animals were protected from disease development (Fig. 4G and 4H).

To evaluate viral replication in tissues, we determined the levels of gRNA and sgRNA by RT-qPCR. We did not identify any differences among study groups in the levels of gRNA and sgRNA in nasal turbinate, lung, in oropharyngeal samples were detected (Fig. 5A and S2C Fig). However, vaccinated animals exhibited a decreasing trend in their nasal turbinate levels of both gRNA and sgRNA over time after challenge (p=0.061; Peto & Peto Left-censored k-sample test) (Fig. 5A and S2C Fig). In addition, the analysis of nasal turbinate samples on day 7 post-challenge showed that vaccinated GSH displayed lower gRNA and sgRNA levels tendency compared with unvaccinated-challenged controls (gRNA: p=0.061; sgRNA: p=0.056; Peto & Peto Left-censored k-sample test) (Fig. 5A and S2C Fig). To confirm RT-qPCR data, the presence of NP was analyzed in nasal turbinate and lung by IHC. NP was not detected in nasal turbinate samples from immunized animals on day 7 (Fig. 5B). These results confirm the decreasing trend observed when gRNA and sgRNA were analyzed over time. Similarly, SARS-CoV-2 replication associated lesions were hardly detected in nasal turbinate samples on day 7 (Fig. 5C). However, despite NP was not detected in lung of vaccinated GSHs on day 7, low levels of tissue lesions were still present (Fig. 5C). No significant differences in tissue damage were observed in lung samples among study groups, probably due to the low number of animals per group.

**Fig. 5.**
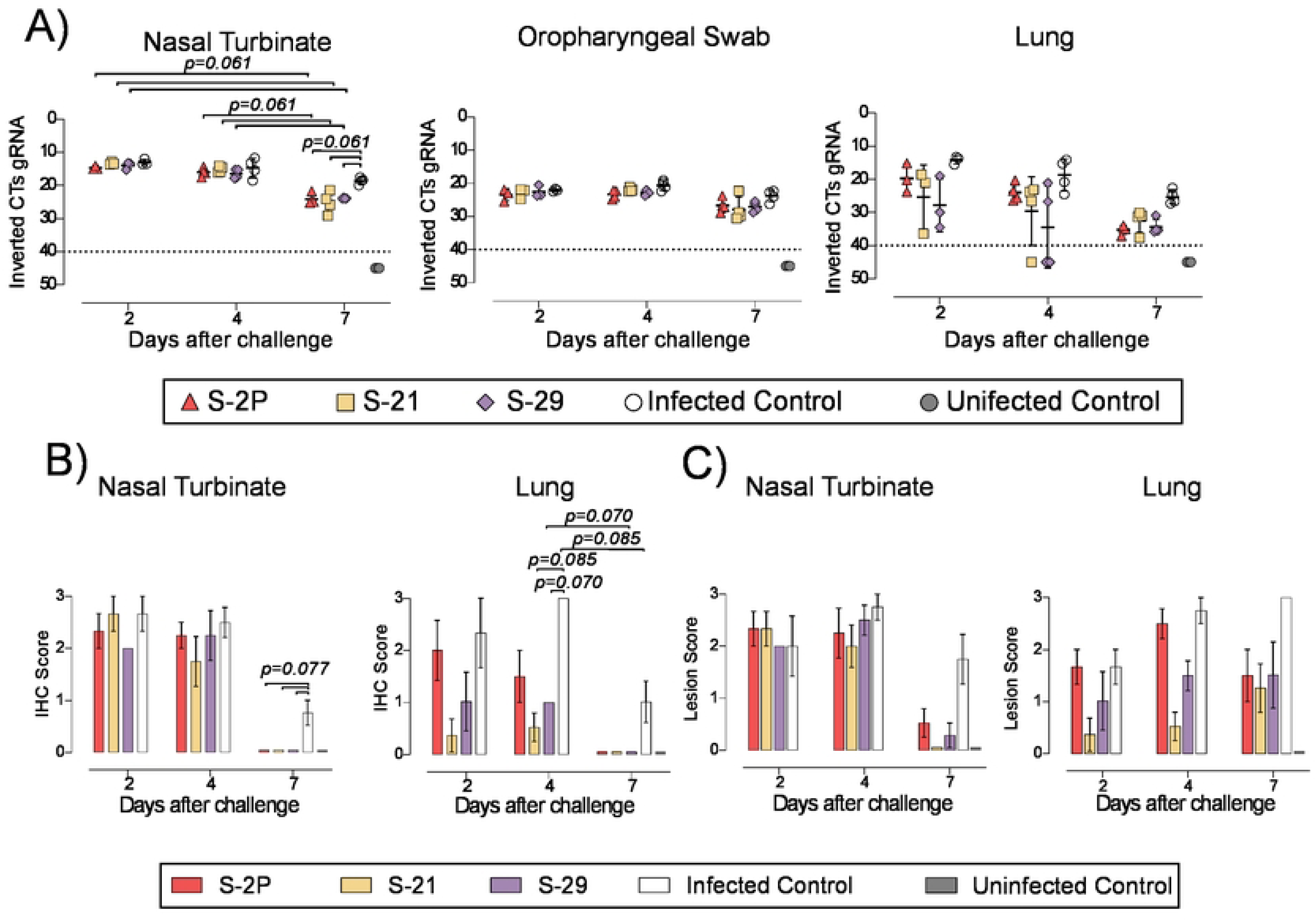
Histopathology and viral loads in tissues from vaccinated GSH after challenge with SARS-CoV-2 Beta variant. SARS-CoV-2 viral loads were analyzed in oropharyngeal swabs, and samples from nasal turbinate and lung of vaccinated and challenged GSH. Virus distribution and tissue damage was analyzed by immunohistochemistry and histopathology, respectively. **A** Levels of SARS-CoV-2 gRNA, expressed as cycles threshold (CTs), in oropharyngeal swabs, nasal turbinate, and lung during infection. Dotted line indicates limit of positivity (40 CTs). Differences among groups were analyzed using Peto & Peto left-censored k-sample test with FDR correction. **B** Histopathological analysis of lung and nasal turbinate by hematoxylin and eosin staining. Lesion score: (0) no lesion, (1) mild lesion, (2) moderate lesion, and (3) severe lesion. **C** Detection of SARS-CoV-2 Nucleocapsid protein in lung and nasal turbinate by IHC. Staining score: (0) no antigen, (1) low, (2) moderate, and (3) high viral antigen. Differences among groups were analyzed using Asymptotic Generalized Pearson Chi-Squared test and FDR.

Overall, these results confirm that all three S-2P, S-21 and S-29 immunogens showed an equivalent immunogenicity and prophylactic activity in GSHs, protecting these animals from the development of severe SARS-CoV-2-induced disease.

## Discussion

The implementation of SARS-CoV-2 vaccines became an inflexion point on the course of the COVID-19 pandemic. However, new SARS-CoV-2 variants have shown partial resistance to the immunity generated by the first generation of COVID-19 vaccines, which were based on the ancestral WH1 sequence[28–30]. Although additional immunizations proved to increase the protection level against new emerging SARS-CoV-2 variants[31,32], this protection remains transient[33]. Particularly, the levels of NAbs elicited against the newest variants (i.e. Omicron and subvariants) wane overtime[34,35], pointing out the importance of developing novel vaccines that increase coverage and duration of immunity. Thus, the adaptation of vaccines to the new variants has shown encouraging results[36–38], and several studies performed in animal models indicate that intranasal immunization may also improve protection[39,40]. Besides these two complementary approaches, S immunogenicity can be enhanced by protein stabilization strategies. Studies performed with the S glycoprotein of MERS and with functional analogous of other viruses have shown that the introduction of mutations that stabilize these proteins in a prefusion state may increase its production and immunogenicity[13,41]. Accordingly, the introduction of two prolines (K986P and V987P) into the S2 subunit of SARS-CoV-2 S was promptly confirmed to enhance its stability and immunogenicity[41], and was successfully implemented in several widely used commercial vaccines (e.g. BNT162b2, mRNA-1273 and Ad26.COV2.S). However, it is still possible to improve the current S-2P strategy since the target recombinant protein is produced with low yield and shows some degree of instability[18]. Initial attempts to stabilize the S in its closed conformation yielded low production, suggesting that the open conformation of the RBD or its intrinsic motility might play a role in protein expression[20]. Recently, Juraszek and colleagues showed that the incorporation of the D614N, A892P, A942P, and V987P mutations were able to stabilize the S glycoprotein in its closed conformation and increase its yield by six-fold compared to the original S-2P protein[19]. Interestingly, Hsieh et al. improved the S stability and production by ten-fold, after introducing four additional proline mutations into the S-2P backbone[18].

Here, we designed and produced a set of S-2P mutated variants whose yield increased between two and five-fold using our new computational pipeline. Unlike proline substitutions, we selected mutations that generated hydrogen bonds, ionic interactions filling hydrophobic pockets, or other well-defined interaction between different chains of the S trimer. We selected two representative S mutants based on their production levels and RBD exposure. Thus, S-21 showed the highest RBD exposure and a moderate increased production, while S29 showed the highest production but a moderate increased RBD exposure compared to S-2P. S-21, S-29 and S-2P were then compared in terms of immunogenicity and capacity to protect against the SARS-CoV-2 Beta variant, one of the most virulent SARS-CoV-2 variants tested in the K18-hACE2 mouse model[24]. To substantiate our results, we used two different animal models: K18-hACE2 transgenic mice and GSHs. The K18-hACE2 is a transgenic mouse model that develops severe disease after SARS-CoV-2 challenge[42,43]. Most mice succumb after viral challenge, mainly due to the infection of the central nervous system[44]. On the other hand, GSHs develop a moderate disease, and animals spontaneously recover two weeks after challenge[26,27]. Our results showed that despite S-2P, S-21, and S-29 showed equivalent immunogenicity and protected both animal models against disease progression, they differed in the degree of protection. The S-29 protein induced an immune response that protected 100% of K18-hACE2 transgenic mice after challenge with the SARS-CoV-2 Beta variant. On the contrary, one mouse in both S-2P and S-21 groups developed severe disease and had to be euthanized on days 8 and 14 after challenge, respectively. Therefore, our results suggested that the S stabilization may impact on the capacity of this protein to induce a protective immune response, particularly against heterologous SARS-CoV-2 variants. According to the *in vivo* protection data, S-29– and S-21-immunized animals showed a faster viral clearance in nasal turbinate than S-2P immunized mice.

In summary, we described a novel set of mutations that stabilized the S glycoprotein, increasing its production *in vitro,* and improving its protective capacity against SARS-CoV-2-induced disease *in vivo*. Despite all these immunogens were based on the original WH1 S sequence, S-29 protein showed 100 % protection against the SARS-CoV-2 Beta variant. The inclusion of these mutations on the next generation of variant-adapted S-based COVID-19 vaccines could enhance the degree of protection to new emerging variants. In addition to an improvement in mucosal vaccine delivery, these advances could significantly contribute to the generation of novel COVID-19 mucosal vaccines that prevent viral infection, irrespectively of the circulating variants. Our results, including our new computational pipeline, may also contribute to the development of novel vaccines for other pathogenic viruses.

## Methods

### 1) Recombinant trimeric S glycoprotein design and modelling

Unsolved secondary structures of the trimer in closed (PDB: 6VXX) and open (PDB: 6VYB) conformations[8] were reconstructed using SwissModel[45]. Then, all possible single mutations in both conformations were modelled using FoldX[46]. For selecting potential variants, two different approaches were used. First, we computed the Gibbs free energy change (ΔΔG_open_) between the WT and the mutant using the open state as a reference. Negative values indicate introduction of stabilization. Second, comparison of the Gibbs free energy changes upon mutation between the closed (ΔΔG_closed_) and open (ΔΔG_open_) conformations (ΔΔG) revealed a set of mutations predicted to strengthen the open conformation (positive values indicate stabilization of the open state). For both approaches, all single mutations predicting beneficial energies (or just slightly neutral/worst values) were addressed by inspecting the three-dimensional models. In this regard, the final selection was based on: i) selection of mutations predicted to increase the stability of the open-conformation using FoldX; ii) selection of mutations predicted to increase the stability of the open-conformation over the closed one using FoldX, iii) selection of mutations creating well-defined intermolecular interactions between the RBD domains (including hydrophobic bonds, π-π interactions and cation-π interactions, ionic bonds, hydrophobic contacts or cavity filling mutations) that would exert a positive impact in the open state or a negative one on the closing motion of the trimer.

### 2) Recombinant protein production and purification

Recombinant trimeric S glycoproteins were designed as previously described by Wrapp[11]. Briefly, the C-terminal end of the S was fused to a T4 foldon trimerization domain in tandem with an 8xhis tag and a strep tag II. The furin cleavage site was removed by mutating it to GSAS. DNA constructs were supplied by GeneART (ThermoFisher scientific) as pcDNA3.4-based plasmids. Proteins were produced by transient transfection using the Expi293 expression system (ThermoFisher Scientific), following the manufacturer instructions. Five days after transfection, cell culture supernatants were harvested and clarified by centrifugation (3000xg for 20 minutes) or using Sartoclear Dynamics® Lab V (Sartorius). Supernatants were then filtered at 0.2 μm using Nalgene Rapid Flow sterile single using vacuum filter units (ThermoFisher Scientific) and were purified using Ni-Sepharose Excel histidine-tagged protein purification resin (Cytiva), concentrated, and buffer exchanged to phosphate buffer saline by ultrafiltration (Merck Millipore). Integrity and purity of purified proteins were analyzed by sodium dodecyl sulfate polyacrylamide gel electrophoresis and Coomassie G-250 staining (ThermoFisher Scientific). Purified proteins were stored at –80°C until use.

### 3) Viral stock preparation

*In vivo* challenge experiments were performed using Cat24 SARS-CoV-2 B.1.531 (Beta) variant isolate (EPI_ISL_1663571)[47,48]. Cat24 was isolated from a nasopharyngeal swab from a COVID-19 affected patient, as previously described[47,48] and subsequently grown and titrated in Vero E6 cells (ATCC CRL-1586). Vero E6 cells were cultured in Dulbecco’s modified Eagle medium (Invitrogen) supplemented with 10% fetal bovine serum (FBS, Invitrogen), 100 U/mL penicillin, and 100 μg/mL streptomycin, (all from Invitrogen).

### 4) In vivo immunization and challenge experiments

K18-hACE2 transgenic mice (B6.Cg-Tg(K18-ACE2)2Prlmn/J; stock #034860; Jackson Laboratories) were maintained by breeding K18-hACE2 hemizygous mice with C57BL/6J mice, following the instructions of Jackson Laboratory (https://www.jax.org/strain/034860). Offspring genotyping was performed according to the protocol 38170: Probe Assay – Tg (K18-ACE2)2Prlmn QPCR version 2.0 (https://www.jax.org/Protocol?stockNumber=034860&protocolID=38170). GSH were purchased from Envigo and maintained by brother/sister mating. Both K18-hACE2 transgenic mice and GSH colonies were established at the Centre for Comparative Medicine and Bioimage (CMCiB). Animal studies were evaluated and approved in advance by the Committee on the Ethics of Animal Experimentation of the IGTP and count with the authorization of Generalitat de Catalunya (Code: 10965 and 11094)

Ninety-one K18-hACE2 mice (balanced female-male ratio, 7-9 weeks old) were distributed in five experimental groups: S-2P (n=21), S-21(n=22), S-29 (n=22), unvaccinated and challenged controls (n=16), and uninfected negative controls (n=10). In the case of GSHs, a total of forty-nine animals were used (balanced female-male ratio, 5-7 weeks old) and distributed in five experimental groups: S-2P (n=11), S-21 (n=11), S-29 (n=11), unvaccinated and challenged controls (n=11), and uninfected negative controls (n=5). Both mice and hamsters from S-2P, S-21, and S-29 groups were immunized with 15 μg of recombinant protein with AddaVax^TM^ (Invivogen) as adjuvant in the hock[49]. Three weeks later, immunized animals were boosted with a second dose of the same formulation. Control animals were primed and boosted with PBS and AddaVax^TM^. Two weeks after boosting, mice were challenged with 1000 TCID_50_ of SARS-CoV-2 (Cat24 isolate) and followed up for 14 days. GSHs were challenged three weeks after boosting with 10000 TCID_50_ of SARS-CoV-2 (Cat24 isolate) and followed up for 7 days. After infection, body weight and clinical signs were monitored daily until the end of the experiment. Six mice for each experimental group, except the uninfected controls, were euthanized on days 3 and 6. The remaining mice were followed up until day 14 post-infection. Three and four hamsters from each experimental group, except the uninfected control group, were euthanized on days 2 and 4, respectively. The remaining GSHs were euthanized on day 7 post infection. In both challenge experiments, uninfected control group was euthanized at the end of the experiment. In addition, any animal showing weight loss higher than 20%, a drastic lack of motility, or a significant reduction of their response to stimuli were euthanized according to the humane endpoints defined in the supervision protocol. Biological samples were collected after euthanasia, including oropharyngeal swab, nasal turbinate, lung, and brain (only in the case of mice) to determine viral loads and perform histopathological analysis. Furthermore, blood samples were collected before each immunization, viral challenge, and under euthanasia. Blood was left at room temperature for two hours for clotting, and serum was collected after centrifugation (10 minutes at 5000xg) and stored at –80°C until use.

### 5) Quantification of anti-S and anti-RBD antibodies by ELISA

An in-house ELISA was developed to evaluate IgG antibodies elicited against the S and RBD glycoproteins in serum samples obtained as described before. Nunc MaxiSorp ELISA plates were coated overnight at 4°C. Half plate was coated with 50 ng/well of antigen diluted in PBS (S or RBD, Sino Biological) and the other half plate was incubated only with PBS. Next day, the whole plate was blocked for two hours at room temperature using blocking buffer [PBS with 1% of bovine serum albumin (BSA, Miltenyi Biotech)]. After that, 50 µL of the appropriate standard or diluted samples were added to each half plate in duplicates and incubated overnight at 4°C. All samples were prepared in blocking buffer. For the mouse standard curve, we used anti-6xHis antibody His.H8 (ThermoFisher Scientific) starting at 1 µg/mL followed by 3-fold dilutions. For GSH standard, a positive GSH serum was used. GSH standard was prepared as seven 1/3 dilution of a stating 1/100 dilution. To reduce inter-assay variability, plates were run in parallel, and each plate contained samples from all experimental groups. The following day, plates were incubated with detection antibodies for 1 hour at room temperature. HRP conjugated (Fab)2 Goat anti-mouse IgG (Fc specific (1/20,000 dilution), or Goat anti-hamster IgG (H-L) (1/20,000 dilution) (all from Jackson ImmunoResearch) were used as secondary antibodies in the mouse and GSH IgG ELISA, respectively. Finally, plates were revealed with o-phenylenediamine dihydrochloride (OPD) (Sigma Aldrich) and stopped using 2N of H_2_SO_4_ (Sigma Aldrich). Optical density (OD) was measured at 492 nm with noise correction at 620 nm. The background OD obtained from the half antigen-free plate was subtracted from the half antigen-coated plate to obtain the specific signal for each sample. Data are showed as arbitrary units (AU/ml) according to the standard used.

### 6) Neutralizing activity of serum samples

Sera neutralizing activity was evaluated as described by Pradenas et al.[50]. HIV reporter pseudoviruses expressing SARS-CoV-2 S glycoprotein and carrying the luciferase gene were produced in Expi293F cells (ThermoFisher Scientific) by co-transfection of the pNL4-3. Luc.R-. E-plasmid (NIH AIDS Reagent Program[51]) and plasmids coding for the following SARS-CoV-2 S glycoproteins lacking the last 19 amino acid in C-terminal: Wuhan (WH1), Beta, Delta, or Omicron variants. VSV-G-pseudotyped pseudoviruses were used as negative controls. Transfections were performed with ExpiFectamine293 Reagent kit (ThermoFisher Scientific). Forty-eight hours later, cell supernatants were harvested, filtered at 0.45 µm and frozen at –80°C until use. Pseudovirus titration was performed using HEK293T cells overexpressing WT human ACE-2 (HEK293T/hACE2) (Integral Molecular, USA).

Serum samples were inactivated at 56°C for 60 minutes before use. Once inactivated, serum samples were serially diluted 1/3 in cell culture medium (RPMI-1640, 10% fetal bovine sera) (range 1/60–1/14,580) and incubated with 200 TCID_50_ of SARS-CoV-2-derived pseudoviruses for 1 hour at 37°C. Then, 1×10^4^ HEK293T/hACE2 cells treated with DEAE-Dextran (Sigma-Aldrich) were added. After 48 hours, plates were read using BriteLite Plus Luciferase reagent (PerkinElmer, USA) in an EnSight Multimode Plate Reader. Neutralizing activity was calculated using a four-parameter logistic equation in Prism 8.4.3 (GraphPad Software, USA) and visualized as normalized ID_50_ (reciprocal dilution inhibiting 50% of the infection).

### 7) Viral load quantification in oropharyngeal swab and tissue samples

After euthanasia, oropharyngeal swab and samples from nasal turbinate, lung, and brain were collected in 1.5 mL tubes with DMEM media supplemented with penicillin (100 U/mL) and streptomycin (100 µg/mL). Tissues were homogenized twice at 25 Hz for 30 seconds using a Tissue Lyser II, and a 1.5 mm Tungsten bead (QIAGEN). After that, samples were centrifuged for 2 minutes at 2000xg and supernatants were collected and processed using the Viral RNA/Pathogen Nucleic Acid Isolation kit and a KingFisher instrument (ThermoFisher Scientific), or an IndiMag pathogen kit (Indical Bioscience) on a Biosprint 96 workstation (QIAGEN), following manufacturer’s instructions.

PCR amplification was based on the 2019-Novel Coronavirus Real-Time RT-PCR Diagnostic Panel guidelines, following the protocol developed by the American Center for Disease Control and Prevention (https://www.fda.gov/media/134922/download). Briefly, 20 μL PCR reaction was set up containing 5 μL of RNA, 1.5 μL of N2 primers and probe (2019-nCov CDC EUA Kit, Integrated DNA Technologies) and 10 μl of GoTaq 1-Step RT-qPCR (Promega, Madison, WI, USA). Thermal cycling was performed at 50°C for 15min for reverse transcription, followed by 95°C for 2 min and then 45 cycles of 95°C for 10 sec, 56°C for 15 sec and 72°C for 30 sec in the Applied Biosystems 7500 or QuantStudio5 Real-Time PCR instruments (ThermoFisher Scientific). For absolute quantification, a standard curve was built using 1/5 serial dilutions of a SARS-CoV2 plasmid (2019-nCoV_N_Positive Control, 200 copies/μL, Integrated DNA Technologies) and run in parallel in all PCR determinations. Triplicates were performed to determine viral load of each sample, which was extrapolated from the standard curve (in copies/mL) and corrected by the corresponding dilution factor. Alternatively, results are shown as Ct or 2-ΔCt.

SARS-CoV-2 subgenomic RNA was quantified as previously described[52] with the following primers (Forward; 5-CGATCTCTTGTAGATCTGTTCTC-3′; Reverse, 5′-ATATTGCAGCAGTACGCACACAA-3′) and probe (5′-FAM-ACACTAGCCATCCTTACTGCGCTTCG-TAMRA-3′). Mouse or GSH *gapdh* gene expression was measured in duplicate for each sample using TaqMan gene expression assay (ThermoFisher Scientific) as amplification control.

### 8) Pathology and immunohistochemistry

Nasal turbinate and lung from mice and GSHs, and additionally brain from mice, were collected after euthanasia and fixed by immersion in 10% buffered formalin and embedded into paraffin. Then, tissue slides were stained with hematoxylin/eosin and examined by optical microscopy to be analyzed histopathologically. Samples were scored semi-quantitatively based on the level of inflammation (0-No lesion; 1-Mild, 2-Moderate, or 3-Severe lesion) as described in [27,44].

The levels of SARS-CoV-2 Nucleoprotein in tissue slides were determined by immunohistochemistry. A rabbit monoclonal antibody 40143-R019 (Sino Biological) at 1:15,000 dilution, and the EnVision®+ System linked to horseradish peroxidase (HRP, Agilent-Dako) and 3,3’-diaminobenzidine (DAB) were used. A semi-quantitative score was used to measure the amount of viral antigen in the analyzed tissues (0-No antigen detection, 1-low, 2-moderate and 3-high amount of antigen) [27,44].

### 9) Statistical analysis

Anti-S and anti-RBD IgG data, as well as neutralizing activity differences among groups at each time point were analyzed using Kruskal-Wallis and Conover’s post-hoc tests with multiple comparison correction by using false discovery rate (FDR). Differences among animals within a particular group along time were analyzed using the Friedman test and Conover’s post-hoc tests for paired data and corrected for multiple comparison by FDR. Krustal Wallis and Dunn’s post-hoc test were used in weight variation in SARS-CoV-2 challenged animals. Severe disease incidence was represented by Kaplan Meier plots and Mantel-Cox test was implemented to calculated statistical differences against uninfected group. To analyze SARS-CoV-2 gRNA and sgRNA data, a Peto & Peto Left-censored k-sample test corrected by FDR was performed. Asymptotic Generalized Pearson Chi-Squared test with FDR correction was applied to histopathology analysis. P values are indicated as follows: * p<0.05, ** p<0.01, *** p<0.001, **** p<0.0001. Statistical analyses were conducted using the R software environment (version 4.1)

## Acknowledgements

This work was supported by Grifols pharmaceutical, the CERCA Program (2017 SGR 252; Generalitat de Catalunya), Direcció General de Recerca i Innovació en Salut (Generalitat de Catalunya) (projects SLD0015 and SLD0016), the Carlos III Health Institute (PI17/01518 and PI18/01332), and the crowdfunding projects “YomeCorono”, BonPreu/Esclat, and Correos. JB is supported by the Health Department of the Catalan Government (Generalitat de Catalunya). CAN, APG and PAR were supported by predoctoral grants from Generalitat de Catalunya and Fons Social Europeu (2020 FI_B_0742; 2022 FI_B_00698 and 2020FI_B2_00138, respectively). EP was supported by a doctoral grant from National Agency for Research and Development of Chile (ANID: 72180406). NI-U is supported by the Spanish Ministry of Science and Innovation (grant PID2020-117145RB-I00), EU HORIZON-HLTH-2021-CORONA-01 (grant 101046118). This study was also supported by CIBER – Consorcio Centro de Investigación Biomédica en Red (CB 2021), Carlos III Health Institute, Ministerio de Ciencia e Innovación and Unión Europea – NextGenerationEU. We would like to thank Foundation Dormeur that support the acquisition of the QuantStudio-5 real time PCR system and Eclipse Ts2R-FL Inverted Research Microscope.

Funders had no role in study design, data analysis, decision to publish, or manuscript preparation.

We thank to the CMCiB’s staff (Sara Capdevila, Jordi Grifols, Rosa Maria Ampudia, Jorge Diaz, Yaiza Rosales and Sergi Sunyé) and the BSL3 IRTA-CReSA staff (Xavier Abad, Ivan Cordon, Anna Pou, Oscar García, Joanna Wiacek, Maria Angeles Osuna, Luís Ribas and Claudia Pereira Sunyé) for their technical assistance with *in vivo* animal studies.

## Author contributions

JC, BC, JS, JB, JVA, NIU, VG and AV: Study conception, design, and funding JC, NPL, CAN, PAR: Manuscript draft preparation CAN, BT, PAR, EAE, MB, NPL, MLR, VU, JR, NR, MP, EP, SM, EB, ERM, GC, FTF, APG, CR, CA, RO, AB, RL, DPZ, JMB, NIU, JB, JVA, VG, JS, and JC: Data acquisition, analysis, and interpretation.

## Competing interests

The present work counted on funding from Grifols pharmaceutical. However, Grifols had no role in study design, data analysis, decision to publish or manuscript preparation. PAR is currently employed by Sanofi, which has no association with any content related to this work. The authors declare no other competing interests. Unrelated to the submitted work, JB and JC are founders and shareholders of AlbaJuna Therapeutics, S.L. BC is founder and shareholder of AlbaJuna Therapeutics, S.L. and AELIX Therapeutics, S.L, and V.G is founder and shareholder of Nostrum Biodiscovery…

Unrelated to the submitted work, NI-U is supported by institutional funding from Pharma Mar, HIPRA, Amassence and Palobiofarma.

## Materials & Correspondence

Correspondence and requests for materials should be addressed to: Jorge Carrillo, IrsiCaixa AIDS Research Institute, Germans Trias i Pujol Hospital, Carretera Canyet s/n, Badalona, 08916, Spain.; Joaquim Segalés, Unitat Mixta d’Investigació IRTA-UAB en Sanitat Animal, Centre de Recerca en Sanitat Animal (CReSA), Campus de la Universitat Autònoma de Barcelona (UAB), 08193 Bellaterra, Barcelona, Spain.; and Victor Guallar, Life Sciences Department, Barcelona Supercomputing Center (BSC), Plaça Eusebi Güell, 1-3, 08034, Barcelona, Spain.

## Figure Legends

**S1 Fig.** Vaccine-induced anti-S IgG responses and levels of tissue sub-genomic RNA in immunized K18-hACE2 transgenic mice after challenge with SARS-CoV-2 B.1.351 (Beta) variant. A Kinetics of anti-S IgG antibodies in serum samples. Red triangles: S-2P group (n=21). Yellow squares: S-21 (n=22). Purple diamond: S-29 (n=22). White circles: unvaccinated-challenge mice (n=16). Grey circles: unvaccinated-uninfected mice (n=10). Groups in each time point were analyzed using Kruskal-Wallis and Conover’s post-hoc tests with multiple comparison correction by FDR. Differences among animals within a particular group along time were analyzed using the Friedman test corrected for multiple comparison using FDR. **B** Kinetics of anti-S and anti-RBD IgG antibodies in serum samples after SARS-CoV-2 B.1.351 challenge. Mean plus standard error of the mean (SEM) is shown. **C** Levels of SARS-CoV-2 subgenomic RNA (represented as inverted Ct) in oropharyngeal swabs, nasal turbinate, lung, and brain after virus challenge. Dotted line indicates limit of detection (40 Cts). Differences among groups were analyzed using Peto & Peto left-censored k-sample test corrected by FDR. * p<0.05, ** p<0.01,

**S2 Fig.** Anti-S IgG responses and levels of sub-genomic RNA in tissues of immunized GSH after challenge with SARS-CoV-2 B.1.351 (Beta) variant. **A** Kinetics of anti-S antibodies in serum samples. Red triangles: S-2P group (n= 11). Yellow squares: S-21 (n=11). Purple diamond: S-29 (n=11). White circles: unvaccinated-challenge mice (n=11). Grey circles: unvaccinated-uninfected mice (n=5). Groups in each time point were analyzed using Kruskal-Wallis and Conover’s post-hoc test with multiple comparison correction by FDR. Differences among animals within a particular group along time were analyzed using the Friedman test corrected for multiple comparison using FDR. **B** Kinetics of anti-S and anti-RBD IgG antibodies in serum samples at days 2, 4 and 6-7 (end point) after SARS-CoV-2 Beta challenge. Mean plus standard error of the means (SEM) are shown. **C** Levels of SARS-CoV-2 subgenomic RNA (represented as inverted Ct) in oropharyngeal swabs, nasal turbinate, and lung after virus challenge. Dotted line indicates limit of positivity (40 Cts). Differences among groups were analyzed using Peto & Peto Left-censored k-sample test corrected by FDR. * p<0.05, ** p<0.01, *** p<0.001.

